# Genome-wide analysis of mobile element insertions in human genomes

**DOI:** 10.1101/2021.01.22.427873

**Authors:** Yiwei Niu, Xueyi Teng, Yirong Shi, Yanyan Li, Yiheng Tang, Peng Zhang, Huaxia Luo, Quan Kang, The Han100K Initiative, Tao Xu, Shunmin He

## Abstract

Mobile element insertions (MEIs) are a major class of structural variants (SVs) and have been linked to many human genetic disorders, including hemophilia, neurofibromatosis, and various cancers. However, human MEI resources from large-scale genome sequencing are still lacking compared to those for SNPs and SVs. Here, we report a comprehensive map of 36,699 non-reference MEIs constructed from 5,675 genomes, comprising 2,998 Chinese samples (∼26.2X, NyuWa) and 2,677 samples from the 1000 Genomes Project (∼7.4X, 1KGP). We discovered that LINE-1 insertions were highly enriched at centromere regions, implying the role of chromosome context in retroelement insertion. After functional annotation, we estimated that MEIs are responsible for about 9.3% of all protein-truncating events per genome. Finally, we built a companion database named HMEID for public use. This resource represents the latest and largest genomewide study on MEIs and will have broad utility for exploration of human MEI findings.

## Introduction

Transposable elements (TEs), also known as transposons or mobile elements, comprise a significant portion in mammalian genomes (Smit 1999; Deininger et al. 2003; Cordaux and Batzer 2009), approximately half of the human genome (Lander et al. 2001). Most TEs are transposition incompetent due to accumulated interior mutations and truncation or various host repression mechanisms (Goodier 2016). In humans, *Alu*, long interspersed nuclear element 1 (L1), SINE-VNTR-*Alu* (SVA), and HERV-K (also known as HML-2) are four families of TEs which are still active and capable of creating new insertions (Mills et al. 2007; Huang et al. 2012), termed mobile element insertions (MEIs). The transposition events have the potential to disrupt normal gene function and alter transcript expression or splicing at the sites of integration, contributing to disease (Payer and Burns 2019). For example, over 120 TE-mediated insertions have been associated with various human genetic diseases, including hemophilia, Dent disease, neurofibromatosis and various cancers (Hancks and Kazazian 2016). Apart from the impact through insertion events, intrinsic sequence properties of TEs endow some MEIs with functional effects on the host (Payer and Burns 2019), making MEIs differ qualitatively from typical forms of SVs like copy number variants (CNVs). Another important question related to MEIs is the integration site preference, which are usually non-random and influenced by various factors such as DNA sequences and chromatin context (Sultana et al. 2017).

However, despite these important functions, integrated resources for polymorphic TEs in human genomes is still lacking (Goerner-Potvin and Bourque 2018), which could offer a large pool of MEIs to explore TE diversity and serve as bedrock for phenotype-variant association studies. And MEIs are not routinely analyzed in most population-scale whole-genome sequencing (WGS) projects (The 1000 Genomes Project Consortium 2015; Wu et al. 2019, 2019; Cao et al. 2020). To date, the largest and most recent population study of MEIs using WGS remains the one conducted by the 1KGP, which included 2,504 genomes across 26 human populations (Sudmant et al. 2015; Gardner et al. 2017). However, the sequencing depth of the 1KGP is low, which may limit the MEI detection sensitivity and accuracy (Rishishwar et al. 2016). In addition, current MEI genetic resources are mainly from European ancestry cohorts, and the lack of Chinese cohort genomic study on MEIs is a critical part of the missing diversity.

In this study, we employed WGS of 5,675 members from newly sequenced Chinese samples and the 1KGP to construct a resource for non-reference MEIs. Although the 1KGP dataset has already been investigated for MEIs (Sudmant et al. 2015; Gardner et al. 2017), we included it here to increase population diversity and build a comprehensive MEI map. The NyuWa dataset has been used to study spectrum of small variant and build reference panel (Zhang et al. 2020), and the MEIs were not explored yet. Combining two cohorts enabled us to systematically analyze the genomic distribution, mutational patterns, and functional impacts of MEIs. From these analyses, we found that L1 MEIs were highly enriched in centromere, and we determined that MEIs represent about 9.3% of all protein-truncating events per individual, emphasizing the importance of detecting MEI routinely in WGS studies. We have built a companion database named HMEID (available at http://bigdata.ibp.ac.cn/HMEID/) for polymorphic MEIs, which could be explored for new insights into MEI biology.

## Results

### A Comprehensive Map of Non-reference Human MEIs

To generate a comprehensive map of MEIs from human genomes, we jointly analyzed two WGS datasets using MELT (Gardner et al. 2017), the low-coverage 1KGP dataset consisting of 2,677 individuals sequenced to ∼7.4X coverage (Sudmant et al. 2015) and the high-coverage NyuWa dataset including 2,998 Chinese samples sequenced to ∼26.2X coverage (Table S1) (Zhang et al. 2020). After site quality filtering, a total of 36,699 non-reference MEIs were kept, including 26,553 *Alu*s, 7,353 L1s, 2,667 SVAs and 126 HERV-Ks (Table 1). Most *Alu* and L1 MEIs were well-supported by split reads (Fig. S1A) and target site duplications (TSDs) (Fig. S1B). Using Hardy-Weinberg equilibrium (HWE) metrics as a rough proxy of genotyping accuracy, we found that about 87% autosomal MEI sites did not violate the HWE, and when restricted to the NyuWa dataset, almost all MEIs (97%) on autosomes had high genotyping accuracy (Fig. S2).

**Table 1.** MEI discovery in this study.

|  | Total sites | Mean sites per donor |  | Standard deviation |  |
| --- | --- | --- | --- | --- | --- |
|  |  | NyuWa | 1KGP | NyuWa | 1KGP |
| *Alu* | 26,553 | 1,035 | 884 | 25.3 | 153 |
| LINE-1 | 7,353 | 145 | 119 | 8.35 | 19.3 |
| SVA | 2,667 | 44.4 | 28.8 | 4.83 | 9.9 |
| HERVK | 126 | 11 | 8.23 | 1.86 | 2.12 |
| Total | 36,699 | 1,236 | 1,040 | 30 | 178 |

On average, we detected 1,236 MEIs with each genome in the NyuWa dataset and 1,040 MEIs in the 1KGP dataset (Table 1), which were expected as increased sequencing depth provides more power for MEI detection (Fig. S1C). The smaller correlation between MEI number and sequencing coverage in the NyuWa dataset than that of the 1KGP dataset reflected that MEI detection sensitivity was close to saturation in ∼30X genomic coverage, consistent with the previous evaluation by the authors of MELT (Gardner et al. 2017). The distribution of MEI numbers per individual, MEI allele frequencies and length estimates largely fit the findings of previous studies (Fig. 1) (Gardner et al. 2017, 2019). About 70.7% MEIs are very rare (allele frequency < 0.1%), with over 30% singletons of all four MEI types (Fig. 1C; Fig. S1D). Since a large proportion of MEIs were individual-specific, we next sought to evaluate MEI discovery by increasing sample size. Through randomly down-sampling to different sizes with 100-sample intervals, we estimated the total MEI variants and the increase of variants at different sample sizes (Fig. S1E-I). As expected, we found that the number of all four MEI types continued to rise with the increasing sample size, but the growth rate decreased. When looking at the subfamilies of MEIs, we found that the distributions of active *Alu* and L1 MEIs were in line with previous observations in humans (Gardner et al. 2017; Bennett et al. 2008; Stewart et al. 2011; Hormozdiari et al. 2013), e.g. *Alu*Ya5 and *Alu*Yb8 were found to be the most abundant two *Alu* subfamilies (Fig. S3), indicating their high retrotransposition activity in modern humans.

**Fig 1.**
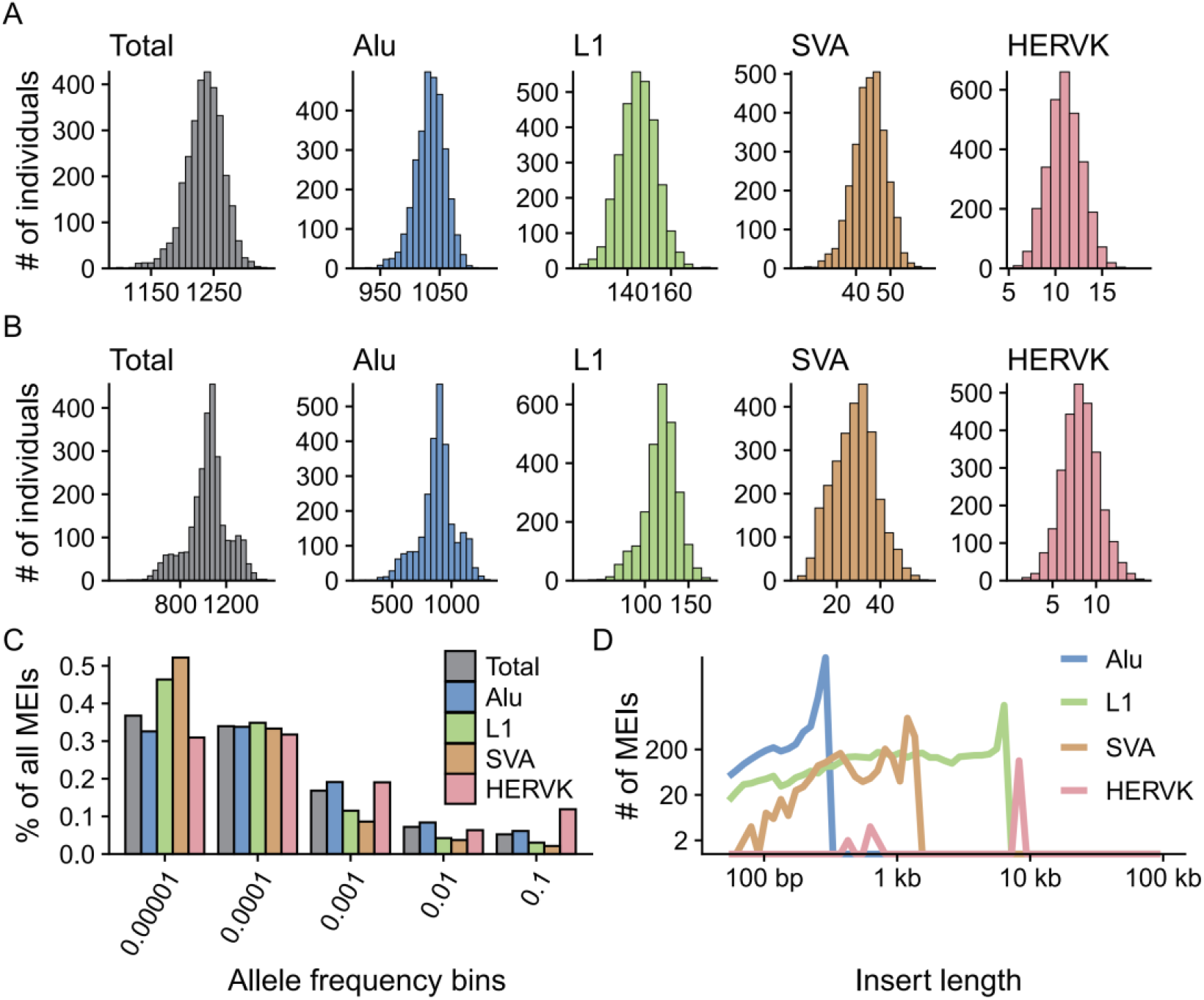
The MEI call set. (**A**) Histograms of the number of MEIs identified per genome in the NyuWa dataset. (**B**) Histograms of the number of MEIs identified per genome in the 1KGP dataset. (**C**) Distribution of allele frequency of MEIs of four types: *Alu*, L1, SVA, and HERVK. “Total” combined the four types of MEIs. (**D**) Distribution of insert size estimated by MELT.

Compared to the previous MEI findings of 1KGP samples (Gardner et al. 2017), the total number of non-reference MEIs we detected has increased 55.4%, with 45.2% and 74.0% increase for *Alu* and L1 insertions respectively (Fig. S4A). In addition, large proportions of MEI calls detected by previous study were repeatedly identified in this study, and the allele frequency for overlapping sites also showed high consistency (Fig. S4B; Pearson’s correlation coefficient = 0.95). Nonetheless, we noticed that many MEIs identified by Gardner *et al*. (Gardner et al. 2017) were missed in our call set. We conjectured that this may be due to differences of software version, reference genome build, and the way how the BAM files were generated etc. To test this, we performed three runs using three sample sets: 1) 100 samples from the 1KGP with reads mapping to the GRCh37 genome build; 2) 100 samples from the 1KGP with reads mapping to the GRCh38 genome build; 3) 100 samples from the 1KGP and 100 samples from the NyuWa, with reads mapping to the GRCh38 genome build. We found that more MEIs could be detected by using the GRCh38 genome build and/or by combining more samples (Table S2). This is also in line with the model used by MELT (Gardner et al. 2017), combining the 1KGP dataset with the high-coverage NyuWa dataset would improve MEI detection sensitivity as well as accuracy, with finer resolution of MEI break points. Collectively, our MEI call set represents a high-quality map of non-reference MEIs for humans.

### Enrichment of Non-reference L1 insertions in Centromeres

It has been long noted that L1s occur preferentially in AT-rich regions but *Alu*s show the opposite trend (Lander et al. 2001). As expected, we also observed this tendency for MEIs (Fig. S5A). In addition, the GC content of flanking DNA for *Alu*s and L1s were lower than background, while SVAs and HERV-Ks prefer DNA sequences with much higher GC content. We next compared the GC composition of rare MEIs (allele frequency < 1%) and common MEIs (allele frequency >= 1%) due to the reported bias shift in GC bias for older and younger short interspersed nuclear elements (SINEs) (Smit 1999; Hormozdiari et al. 2013; Medstrand et al. 2002; Waterson et al. 2005). Significant difference was only observed for HERV-K: rare HERV-K insertions occurred in much higher density at GC-rich regions (Fig. S5B). We did not observe marked bias for *Alu*s and SVAs, likely because most insertions we identified were already fixed in population.

We next sought to investigate the distribution of MEIs throughout the genome, like previously Collins *et al*. had done for common SVs (Collins et al. 2020). Interestingly, L1s were predominantly enriched at centromeric regions, whereas SVAs and HERV-Ks were enriched at telomeres (Fig. 2 A and B; Fig. S6). For comparison, similar analysis was applied to TEs in the reference genome, but no such patterns for L1s were found (Fig. S7B). Even in the latest telomere-to-telomere assembly of the human X chromosome, only a single L1 insertion was detected at the centromere region (Miga et al. 2020). When restricted to singleton L1 MEIs, we could still detect the enrichment in centromeres (Fig. 2C). Importantly, this finding was well-supported by non-reference L1s from euL1db (Fig. 2D) (Mir et al. 2015), which curated human polymorphic L1s from 32 different studies. Considering the reduced detection power of short-read WGS in repetitive regions, the enrichment of L1 insertions at centromeric regions could be still underestimated. The enrichment of non-reference L1 insertions at centromeric DNA could be partly attributed to lower GC content, as centromeres contain massive AT-rich alpha satellites (Manuelidis and Wu 1978). Also, active TEs have been found in neocentromere regions, and may contribute to centromere ontogenesis (Klein and O’Neill 2018; Contreras-Galindo et al. 2013; Zahn et al. 2015). The reasons for the dramatic enrichment of L1s in centromere regions are intriguing and further studies are needed in the future.

**Fig 2.**
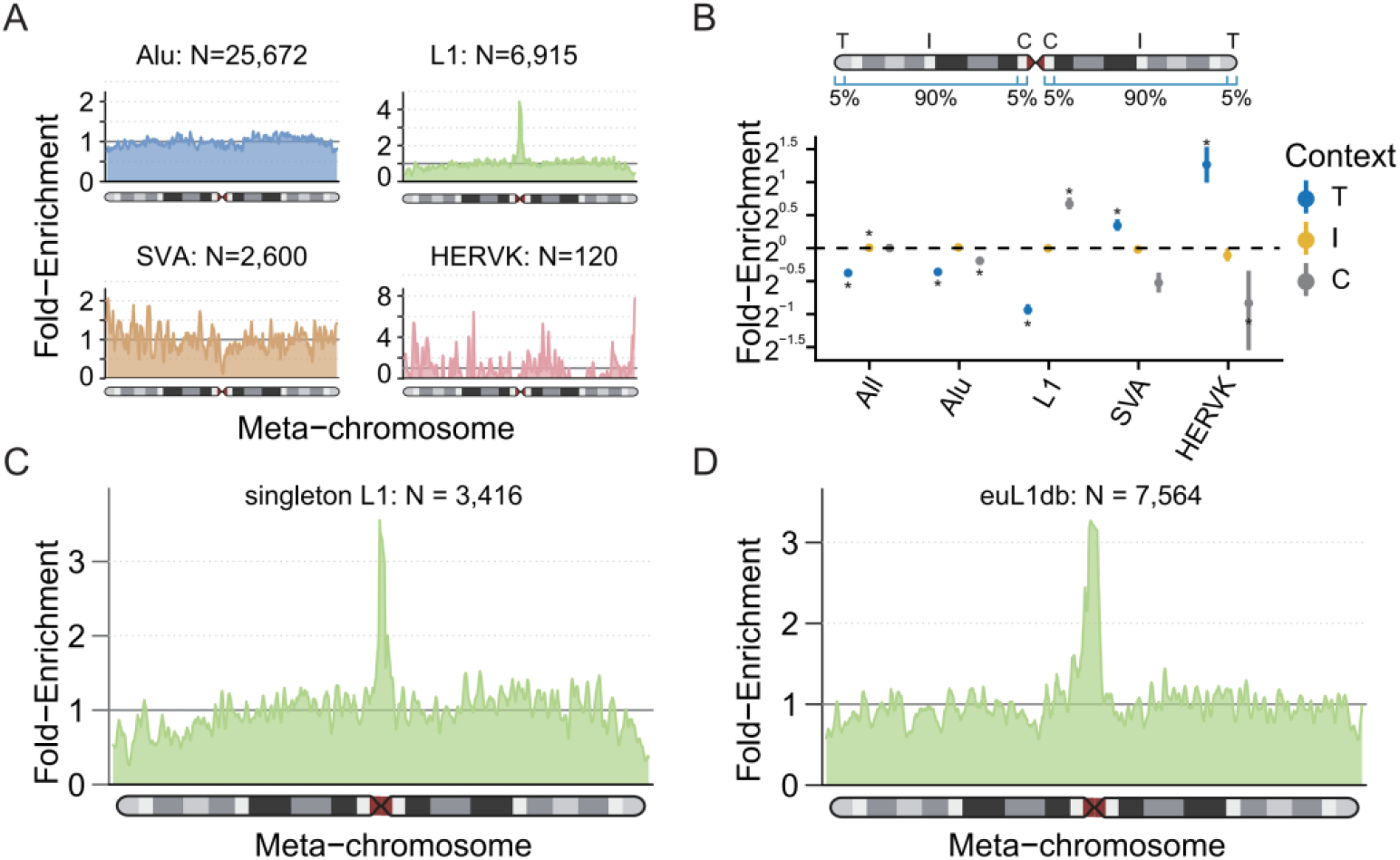
Chromosome-level Distribution of MEI Density. (**A**) Smoothed enrichment of different types of MEIs ascertained in this study. The values were calculated per 100 kb window across the average of all autosomes and normalized by the length of chromosome arms (as “meta-chromosome”). (**B**) Enrichment of MEIs by class and chromosomal context. The dots are the mean values and point ranges represent 95% confidence intervals (CIs). P-values were computed using a two-sided t-test and adjusted using the Bonferroni method. *, p ≤ 0.05. C, centromeric; I, interstitial; T, telomeric. The way to compute the chromosomal enrichment and to represent data was from the gnomAD SV paper (Collins et al. 2020). (**C**) Smoothed enrichment of singleton L1s (L1 MEIs found in single genome) ascertained in this study. (**D**) Smoothed enrichment of non-reference L1s from euL1db database (Mir et al. 2015).

### Strong Correlations between MEI Diversity and SNP Heterozygosity

Since mutations are ultimate sources of genetic innovation and significant causes of human birth defects and diseases, knowledge of mutation rate is a general population genetics question (Kumar and Subramanian 2002; Feusier et al. 2019). Here we employed the commonly-used Waterson’s estimator (Watterson 1975) of Θ to estimate the mutation rate of each MEI type and found that mutation rates varied markedly by MEI class (Table S3). Since MEI detection and genotyping power is profoundly influenced by sample coverage (Gardner et al. 2017), we conducted the analysis separately for the NyuWa and the 1KGP datasets. The resulting calculation provided very close estimates of between 3.217×10^−11^ (NyuWa) and 2.928×10^−11^ (1KGP) de novo MEIs per bp per generation (μ), or roughly one new MEIs genome-wide every 11-16 live births, which is largely concordant with prior reports (Sudmant et al. 2015; Gardner et al. 2019).

The availability of SNP genotyping (both the NyuWa and the 1KGP dataset) for the same samples given us an opportunity to investigate the correlation between MEI diversity and SNP heterozygosity for each population. SNP heterozygosity was computed as the ratio of heterozygous SNPs across the individual’s genome (Prado-Martinez et al. 2013) and was compared to the average MEI differences between samples in a given population (Hedges et al. 2004). The diversity for all types of MEIs showed strong correlation with SNP heterozygosity (R^2^: 0.64∼0.95), with African populations showing the highest MEI diversity and SNP heterozygosity (Fig. 3) — consistent with previous study (Stewart et al. 2011).

**Fig 3.**
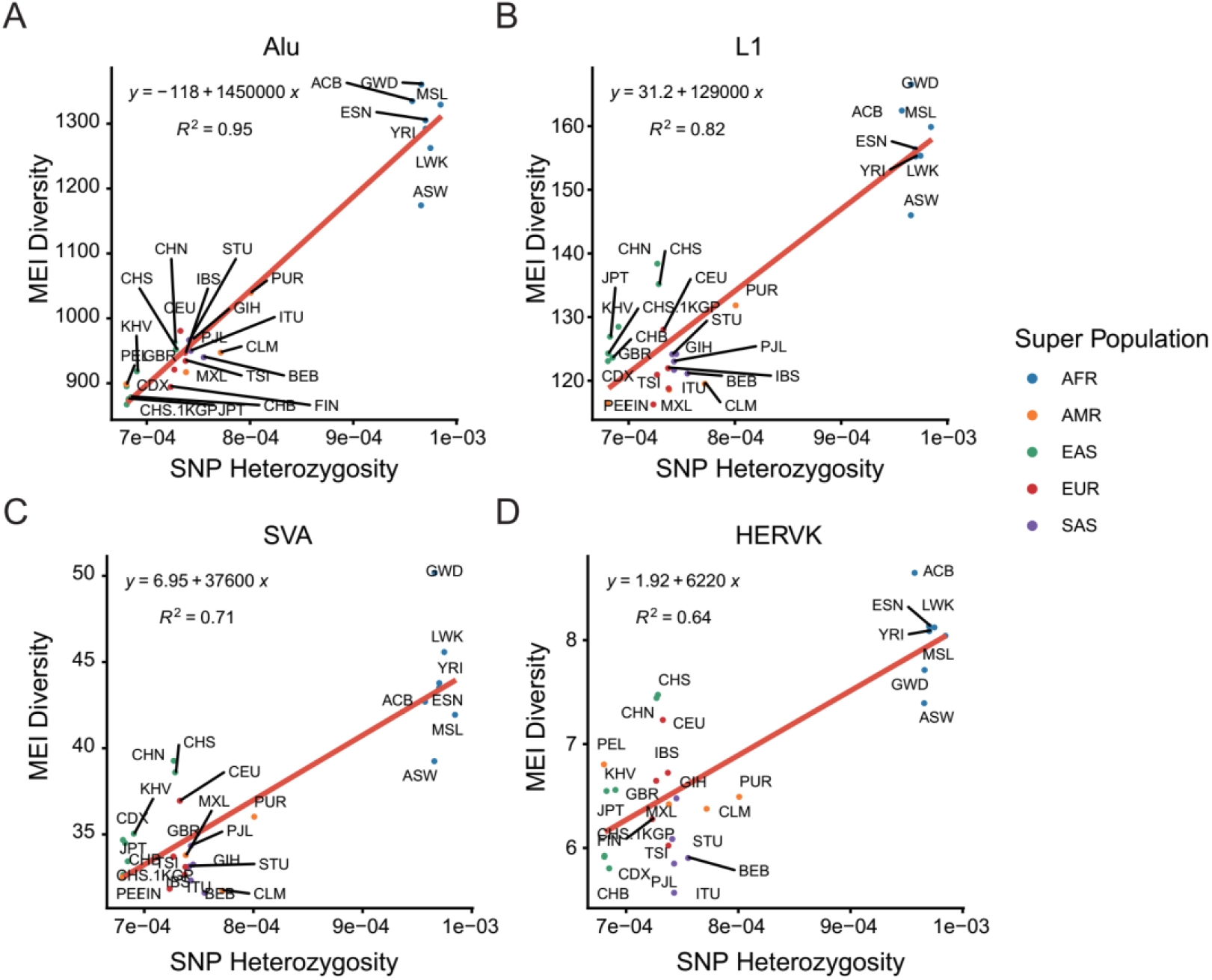
Correlation between SNP heterozygosity and MEI diversity. SNP heterozygosities and diversity of (**A**) *Alu* MEIs, (**B**) L1 MEIs, (**C**) SVA MEIs and (**D**) HERV-K MEIs were compared in different populations. SNP heterozygosity was computed as the ratio of heterozygous SNPs across the individual’s genome and MEI diversity was computed as the average allele difference in each population. Points were colored by super populations. AFR, African super population; AMR, American super population; EAS, East Asian super population; EUR, European super population; SAS, South Asian super population.

### MEI Functional Properties

Via the local impacts by transposition events or more global post-insertion influence (Klein and O’Neill 2018), MEIs can disrupt normal gene functions and be disease-causing (Payer and Burns 2019; Hancks and Kazazian 2016). In principle, any MEIs can result in predicted loss-of-function (pLoF) by altering open-reading frames. To assess the functional impacts of MEIs, we annotated the MEI calls using Variant Effect Predictor (VEP) and BEDtools (see Methods). The vast majority (82.7%) of detected MEIs was in intergenic and intronic regions, while only ∼2.7% MEIs impacted the coding sequences (CDS) (Fig. 4A). Varying enrichment levels on different genomic features were observed for different MEI types (Fig. 4B). For example, L1, SVA and HERV-K MEIs were significantly depleted in CDS and non-coding gene exons; L1 MEIs were enriched in coding introns and gene flanking regions; SVA and HERV-K sites were enriched in intergenic and non-coding introns. Focusing on protein-truncating variants (PTVs), each genome contained a mean of 24.8 MEIs (12.6 *Alu*, 7.4 L1, 1.3 SVA and 2.4 HERV-K) directly disrupting CDS, including 1.1 rare pLoF MEIs (allele frequency < 1%) (Fig. 4C; Table S4). By comparison, Karczewski *et al*. estimated 98.9 pLoF short variants (SNVs and InDels) per genome (Karczewski et al. 2020), and Collins *et al*. observed 144.3 pLOF SVs per genome (Collins et al. 2020). We thus estimated that MEIs account for about 9.3% (24.8/268) of all PTVs, among small variants and large SVs in each human genome.

**Fig 4.**
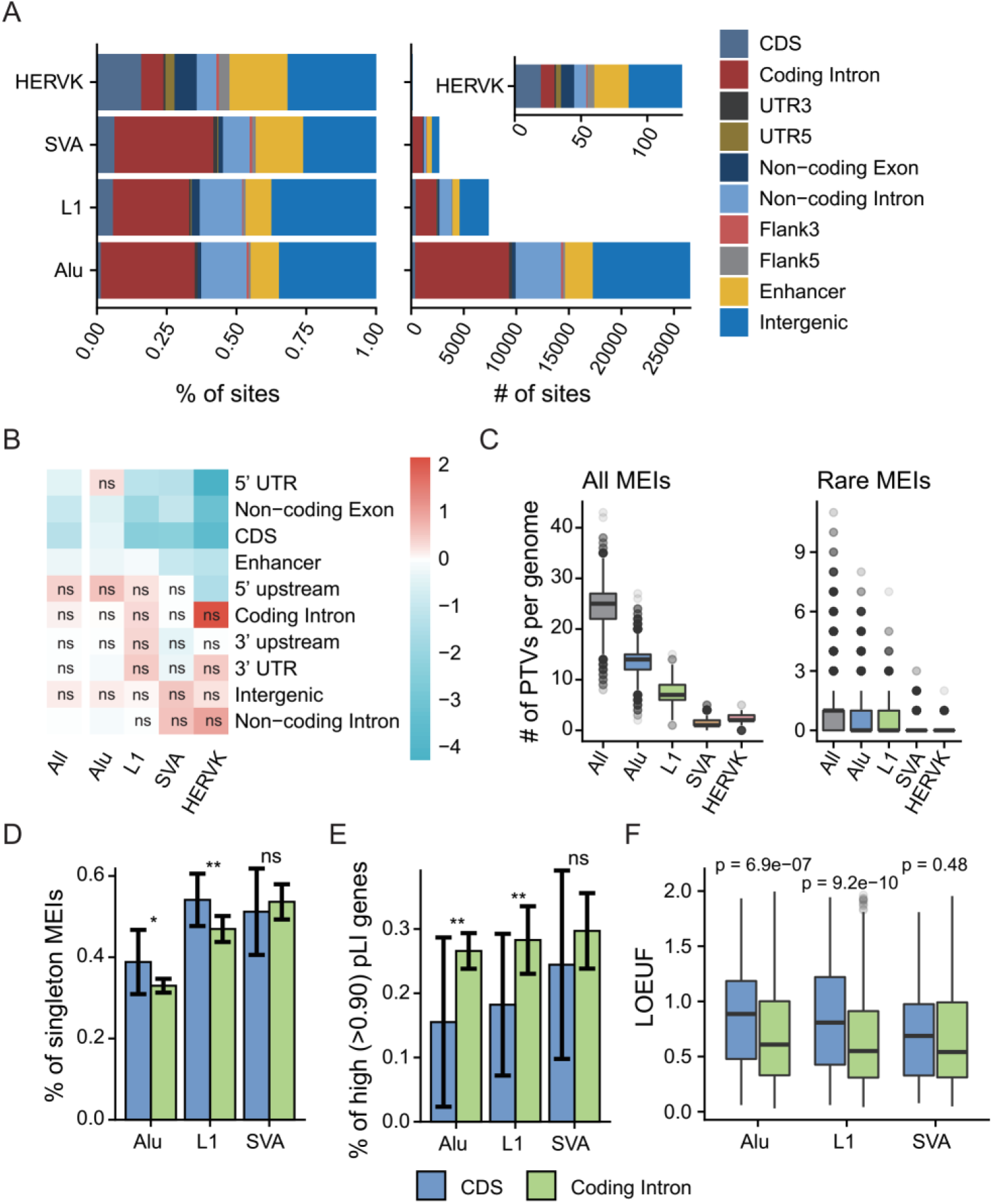
MEI functional properties. (**A**) Predicted functional consequences for each type of MEI: (left) cumulative proportion, and (right) cumulative number. (**B**) Log2 fold enrichment of the MEI call set compared against the MEIs permutated. The permutation test was repeated 1000 times, and empirical p-values were commutated together with the enrichment values. The enrichment values were scaled row-wise. ns, not significant (p-value > 0.05). (**C**) Box plots of counts of predicted PTVs by MEI: (left) all the MEIs identified in this study, and (right) rare MEIs (allele frequency < 1%) in this study. (**D**) Proportions of singleton MEIs in CDS and coding introns for *Alu*, L1 and SVA. Error bars indicate 95% CIs based on population proportion. P-values were computed using chi-squared test. (**E**) Proportions of high pLI genes (pLI > 0.9) for genes with MEIs in the CDS and genes with MEIs intron regions. Error bars represent 95% CIs based on population proportion. P-values were computed using chi-squared test. (**F**) Box plots of LOEUF scores of genes with MEIs in the CDS and genes with MEIs in their introns. Wilcoxon rank sum test was used to compute p-values. Figure D-F used the same legend beneath. ns, p ≥0.05; *, p < 0.05; **, p < 0.01.

Examining the degree to which evolutionary forces acting on coding MEI loci is important to understand the relationships between MEI variation and coding genes. Here we used three different metrics to investigate selective constraints: 1) the proportion of singleton variants (variants observed in only one individual), an established proxy for selection strengths (Lek et al. 2016); 2) the proportion of MEIs in genes with high probability of loss-of-function intolerance (pLI) (Lek et al. 2016); 3) the loss-of-function observed/expected upper bound fraction (LOEUF) of MEI-containing coding genes, where higher LOEUF scores suggest a relatively higher tolerance to inactivation for a given gene (Karczewski et al. 2020). HERV-KMEI was not included in this analysis due to the relatively small number found in coding genes.

Higher singleton proportions for *Alu* and L1 MEIs were found in CDS than that of introns (Fig. 4D; χ2 p < 0.05), while we did not find a statistically significant bias for SVA MEIs, though there were 166 and 949 SVA insertions found in CDS and coding introns, respectively. Likewise, lower proportions of *Alu*/L1 MEIs detected in genes with high pLI score (> 0.9) were found in CDS than that of intronic regions (Fig. 4E; χ2 p < 0.05). Observations from the perspective of enclosing genes fit these results: higher LOEUF score were found for genes with *Alu*/L1 MEIs (Fig. 4F, Wilcoxon p < 0.05). Our results sustained and expanded previous findings on human exome data (Gardner et al. 2019), in which Gardner *et al*. reported that exonic MEIs were under purifying selection.

Although researchers have long noted that most of reference LTR elements and L1s in gene introns are in the antisense orientation with respect to the host genes (Smit 1999; Medstrand et al. 2002), possibly due to ill effects on transcript processing of sense-oriented elements (van de Lagemaat et al. 2006; Zhang et al. 2011), there are no established conclusions about the orientation tendency of non-reference MEIs (Gardner et al. 2019; Hormozdiari et al. 2013). Our large collection of MEIs found in genes allowed us to closely examine the strand bias of different MEIs. Although a bias for *Alu*, L1 MEIs and SVA MEIs to be in an antisense orientation when found within genes was observed (Hormozdiari et al. 2013), we did not find a statistically significant bias for L1 insertions (Fig. S8A). Conversely, *Alu*s were found to have strong strand bias when being inserted into protein-coding genes, non-coding genes, protein-coding introns, and non-coding introns (Fig. S8; χ2 p < 0.05). For SVA MEIs, protein-coding genes, protein-coding exons, and protein-coding introns were regions where insertion orientation biases were detected (Fig. S8; χ2 p < 0.05). Considering that *Alu* and SVA elements are non-autonomous TEs that are trans-mobilized by the L1 retrotransposition machinery (Dewannieux et al. 2003; Raiz et al. 2012), there may be some post-insertion selection forces on *Alu*/SVA elements which influence these patterns (Sultana et al. 2017). The genes themselves which had MEIs in sense or antisense strand in introns did not show clear differences in terms of selective constraints, by comparing the LOEUF scores of these two kinds genes (Fig. S8F). In addition, no significant orientation tendency against the neighboring genes were detected when MEIs were in gene upstream regions (Fig. S8I).

*Alu* MEIs have been found to be enriched in regions of genome associated with human disease risk, suggesting their potential effects on common diseases (Payer and Burns 2019; Payer et al. 2017). To identify MEIs potentially associated with human trait or disease, we mapped MEIs to regions in linkage disequilibrium (LD) with trait- or disease-associated loci identified by genome-wide association study (GWAS) (P < 10^−;8^) (Buniello et al. 2019). We found that 6,457 (about 17.6%) of the MEIs (17.5% for *Alu*, 15.3% for L1, 24.4% for SVA, and 16.6% for HERV-K) were in these regions that tagged by at least by one GWAS SNP (Table S5), with allele frequency of 738 MEIs over 1%, suggesting the remarkable potential for MEIs to contribute in disease and the utility of our MEI set in future phenotype-variant association studies.

### L1 3’ Transduction and 5’ Inversion

Some L1 elements can bring a 3’ readthrough transcript to the offspring insert site, which is called 3’ transduction (Goodier et al. 2000). These L1 elements are usually near a strong Poly(A) sequence. Transcription of these L1 elements is not terminated by the original weak Poly(A) of the L1 element but by the stronger poly(A) sequence downstream. With the flanking sequences downstream L1 elements, we extracted the correspondence between L1s in different genomic positions. Totally, 446 offspring MEIs derived from 57 source MEIs were identified in our samples. These MEI relationships are both interchromosomal and intrachromosomal (Fig. 5A). Compared with L1 transduction source sites identified by 1KGP study (Gardner et al. 2017), we found most of the sites were overlapped (Fig. 5B). Among these sites, 2 of the 3 most active source sites (chr6:13190802, chr1:118858380) were also found in this study, while the site *L1RE3* (chr2: 155671336) is in a low complexity region and was filtered in the site filtering. Most of the sources transducts less than 20 offspring whereas site chrX:11713279 has 186 offspring (41% of all offspring detected). Source and offspring MEIs were distributed into families and population frequency was calculated (Fig. 5C and D). Most transduction classes were EAS specific. Comparing frequencies among subpopulations of EAS, we noticed 14 transduction classes only detected in Chinese people. Inside these classes, 5 classes only appear in samples of Northern Han Chinese (CHB, CHN) and 4 classes only appear in Southern Han Chinese (CHS, CHS.1KGP) (Table S6).

**Fig 5.**
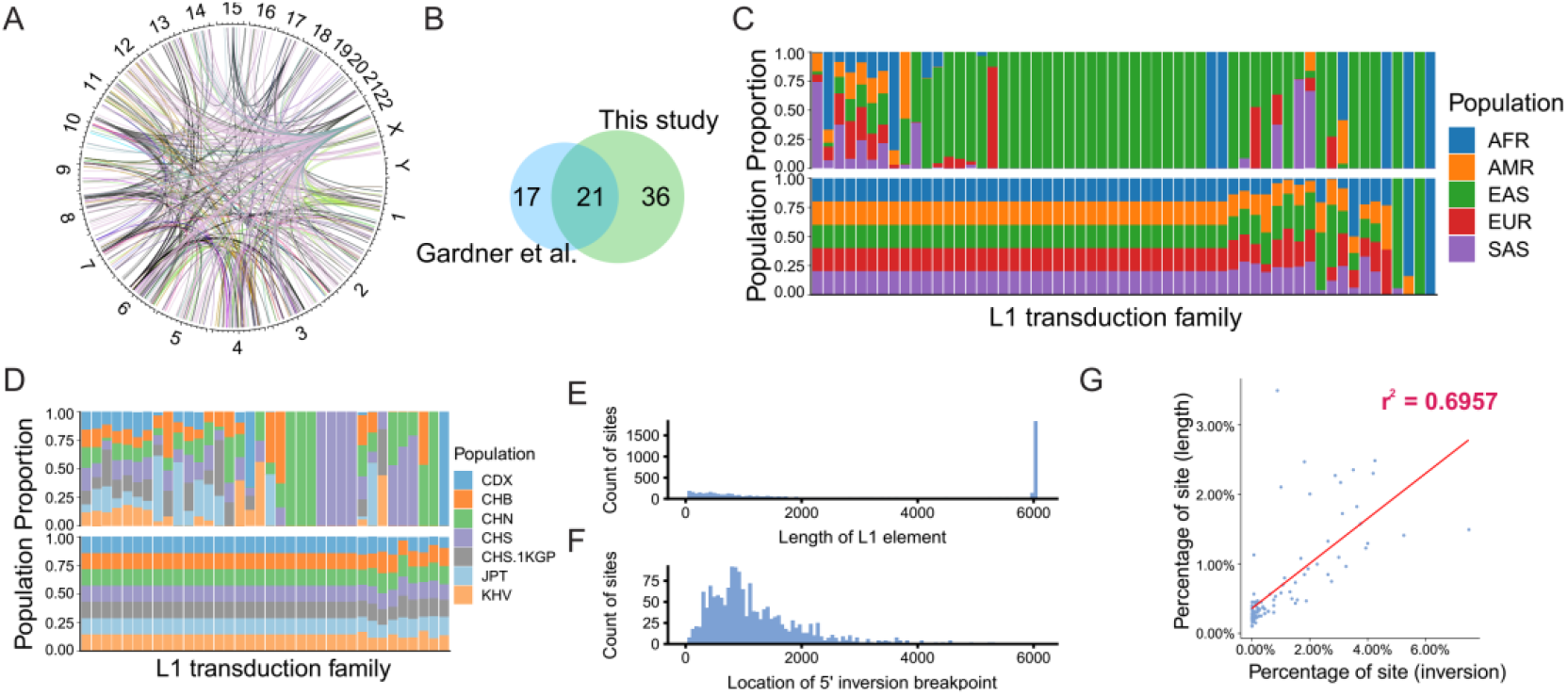
L1 3’ Transduction and 5’ Inversion. (**A**) 3’ transduction source-offspring relations across the whole genome. (**B**) Venn plot of 3’ transduction sources found by our study and the 1KGP study (Gardner et al. 2017). (**C**) Source (bottom) and offspring (top) element frequencies in super populations. AFR, African super population; AMR, American super population; EAS, East Asian super population; EUR, European super population; SAS, South Asian super population. (**D**) Source (bottom) and offspring (top) element frequencies in Asian subpopulations. CDX, Chinese Dai in Xishuangbanna; CHB, Han Chinese in Beijing; CHN, Northern Han Chinese, China; CHS, Southern Han Chinese; CHS.1KGP, Southern Han Chinese from the 1KGP; JPT, Japanese in Tokyo; KHV, Kinh in Ho Chi Minh City. (**E**) L1 length distribution within our call set. The length was estimated by MELT. (**F**) 5’ inversion position distribution among all inverted sites. (**G**) Correlation plot between the distributions shown in (E) and (F). Full length L1 element was excluded in this comparison.

5’ end of the L1 sequence can be inverted during insertion (Ostertag and Kazazian 2001). We extracted the 5’ inversion information from the MELT result, and 1,606 L1 insertions were detected with a 5’ inversion end. The nearest distance from the 5’ inversion site to the 3’ end of the L1 insertion is 602 bp, which is consistent with the 1KGP study (590 bp) (Gardner et al. 2017). It seems that inversion does not occur in the first ∼600 bp from the 3’ end, which may indicate that the inversion process requires at least ∼600 bp DNA sequence. In the previous study, the distribution of the 5’ inversion positions highly correlated with the distribution of L1 MEI lengths. MEIs in our study also showed this trend (R^2^ = 0.696; Fig. 5E-G). We next calculated the percentage of 5’ end inverted MEIs within each 3’ transduction offspring class. The inversion rate across different classes varied and did not correlate with the class size (Table S7). For the biggest class which derived from chrX:11713279, only 25.3% of the offspring had 5’ inversion while a class which only includes 15 offspring had a 40% inversion rate.

### A Database for Polymorphic MEIs

Currently, resources for polymorphic TE findings in human genomes are in high demand (Goerner-Potvin and Bourque 2018). There were two dedicated databases for polymorphic human MEIs: dbRIP (Wang et al. 2006) and euL1db (Mir et al. 2015). However, the former had not been updated since 2012 and the latter was only for human-specific L1 insertions. To fill this gap, we have designed a companion database named HMEID to archive MEIs identified in this study, and to comprehensively catalog the variants on allele frequencies in the NyuWa dataset and the 1KGP dataset. Besides, variant quality metrics and functional annotations are also presented. Compared to dbRIP, HMEID contained more MEIs; the number of L1 insertions in HMEID was comparable with that of euL1db (Fig. 6). Importantly, HMEID contained MEIs detected from samples of Han Chinese, which is the largest ethnic group in the world. We anticipated that this resource would facilitate the exploration of TE polymorphisms and benefit future researches on TEs as well as human genetics.

**Fig 6.**
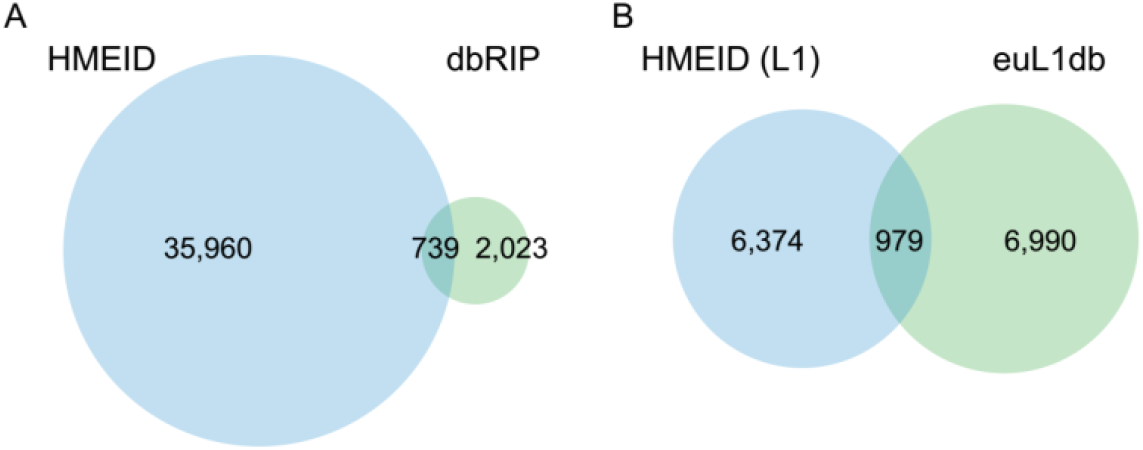
Comparing HMEID with other MEI Databases. (**A**) Comparison the MEI set in the HMEID with that of from the dbRIP database (Wang et al. 2006). (**B**) Comparison the L1 MEIs in the HMEID with non-reference L1s from the euL1db database (Mir et al. 2015).

## Discussion

MEIs, an endogenous and ongoing source of genetic variation, have not been investigated in many population-scale WGS projects. Here we leveraged 5,675 genomes from the NyuWa (Zhang et al. 2020) and the 1KGP (The 1000 Genomes Project Consortium 2015) dataset to identify non-reference MEIs. After describing the frequency spectrum of variants, we focused on the insertion site preference and functional impacts of MEIs. We provided an important resource of non-reference MEIs in humans.

We identified 36,699 non-reference MEIs for four types of TEs and determined that individuals harbour a mean of over 1,000 non-reference MEIs, mostly contributed by *Alu* insertions. In line with previous reports (Gardner et al. 2017, 2019; Stewart et al. 2011), most MEIs were rare and individual-specific, which was also observed for SNVs (The 1000 Genomes Project Consortium 2015) and SVs (Collins et al. 2020). With the newly sequenced 2,998 genomes from China, this study established a large-scale MEI resource for the genetics of Chinese as well as East Asians. Comparing to the previous study conducted by the 1KGP (Gardner et al. 2017), the number of MEIs detected by us has increased about 55%, representing what is to our knowledge the most comprehensive set of human non-reference MEIs.

We found that non-reference MEIs have non-random distributions along chromosomes, implicating the role of chromosome context in TE insertion. Of note, we found that non-reference L1 MEIs were drastically enriched in centromere regions, which was also supported by independent data from the euL1db (Mir et al. 2015). The genomic distribution of TEs is a result from insertion site preference and post-insertion selection on the host (Sultana et al. 2017). On the one hand, human centromeres are full of AT-rich alpha satellites (Manuelidis and Wu 1978), which could confer insertion preference for L1s, since the target specificity of L1 insertion machinery is TTTT/A (Feng et al. 1996). Certain centromeric histones and other centromeric proteins may also serve as preferred targets for TEs, as suggested by a study in maize (Schneider et al. 2016). Additionally, studies on HIV integration into the host genome implied that proximity to the nuclear periphery of centromere may facilitate TE targeting (Lelek et al. 2015; Marini et al. 2015). On the other hand, incorporation of L1s may facilitate the recurring evolutionary novelty of centromeres (Klein and O’Neill 2018). In support of this, Chueh *et al*. reported that RNA transcripts from a full-length L1 are the essential structural and functional components in the regulation of a human neocentromere (Chueh et al. 2009). Evidences were also found in the tammar wallaby (*Macropus eugenii*), where dramatic enrichment of L1s and endogenous retroviruses was found in a latent centromere site (Longo et al. 2009), and *Equus caballus*, where evolutionarily new centromeres locate in LINE- and AT-rich regions (Nergadze et al. 2018). In addition to centromere ontogenesis, a LINE-like element (G2/jockey3) contributes directly to the organization and function of centromeres of *D. melanogaster* (Chang et al. 2019). This is also likely true for the non-reference SVA, for which we found an enrichment in telomeres, as TEs were found to be essential in maintaining the telomere length homeostasis in insects (Pardue and DeBaryshe 2011). However, another plausible explanation for both the enrichment of non-reference L1 MEIs in centromere and non-reference SVA MEIs in telomere is that these regions contain few protein-coding genes, limiting insertional mutagenesis by TEs (Sultana et al. 2017). The reasons for this phenomenon are fascinating, and our study post an important question about the relationship between TEs and centromeres.

Knowing the functional impact of MEIs is fundamental to our understanding the impact of MEI with respect to human disease or trait and evolution (Goerner-Potvin and Bourque 2018). We have estimated that MEIs accounted for about 9.3% of all protein-truncating variants per genome, among small variants (Karczewski et al. 2020) and SVs (Collins et al. 2020). Our estimation was much higher than that determined by whole exome sequencing data (Gardner et al. 2019), possibly due to the limitation of exome baits. We found that a significant portion of polymorphic MEIs mapping to loci implicated in trait/disease association by GWAS, as increasingly recognized by recent studies (Payer et al. 2017; Wang et al. 2017). While previous GWAS have mainly focused on small variants (Visscher et al. 2017), future association studies should consider and evaluate the effects of MEIs in common disease. We anticipate that the HMEID will serve as a basis for such studies.

Our study is limited in that only one tool was used to identify MEIs. Though the overall performance of MELT outperformed existing MEI discovery tools (Gardner et al. 2017) and it has been successfully used in several large-scale studies (Gardner et al. 2017, 2019; Feusier et al. 2019; Werling et al. 2018; Torene et al. 2020), but the detection power could be compromised by modest sequencing depth and incompetence in complex genomic regions of short-read WGS *etc*. In addition, the overall genotyping accuracy by MELT v2 was 87.95% for non-reference *Alu*s (not excluding MEIs in low complexity regions), when compared with PCR generated genotypes (Goubert et al. 2020). As such, we have tried to ensure the site quality by strict filtering. In the future, we would consider combining different MEI identification and genotyping tools to improve the quality, which has been proved useful in previous reports (Ewing 2015; Goerner-Potvin and Bourque 2018; Rishishwar et al. 2016; Feusier et al. 2019).

Also, long-read WGS is promising in detecting MEIs, especially for genomic regions refractory to approaches using short-read sequencing technologies (Audano et al. 2019; Chaisson et al. 2019; Zhou et al. 2020). Another limitation of our MEI dataset is that reference MEIs (MEIs detected as deletions) were not included yet, for which the detection is underway and the results would be integrated into the HMEID for public use.

## Methods

### Experimental design

Data in this study were from two sources: low-coverage (∼7.4X) WGS samples from the 1KGP (The 1000 Genomes Project Consortium 2015) and high-coverage (∼26.2X) WGS samples from the NyuWa dataset (Zhang et al. 2020). For the 1KGP dataset, CRAM-format files of 2,691 individuals were downloaded from http://ftp.1000genomes.ebi.ac.uk/vol1/ftp/data_collections/1000_genomes_project/, which were aligned to human genome build GRCh38 (Lowy-Gallego et al. 2018). The CRAM files were then converted to BAMs using SAMtools v1.9 (Li et al. 2009). The NyuWa dataset contained 2,999 individuals including diabetes and control samples collected from different provinces in China (Zhang et al. 2020), and this cohort was sequenced using the Illumina platform. The processing from raw FASTQs to BAMs was according to the GATK Best Practices Workflows germline short variant discovery pipeline (Poplin et al. 2018), as described in (Zhang et al. 2020). The median depth of the NyuWa samples after genome alignment (GRCh38 human genome build) and removal of PCR duplicates was about 26.2X.

### Generation of MEI call set

MELT v2.15 (Gardner et al. 2017) was run with default parameters using “SPLIT” mode to identify non-reference MEIs, which detects a wide range of non-reference *Alu*, L1, SVA and HERV-K insertions. To get the BAM coverage for MELT analysis, we used goleft v0.1.8 (https://github.com/brentp/goleft) “covstats” function to estimate the genomic coverage for each sample. After initial generation of a unified VCF file by MELT “MakeVCF” function, variants that did not pass the following criteria were filtered to get a high-quality MEI call set: 1) not in low complexity regions; 2) be genotyped in greater than 25.0% of individuals; 3) split reads > 2; 4) MELT ASSESS score > 3; and 5) VCF FILTER column be PASS. 2,998 of 2,999 samples in NyuWa and 2,677 of 2,691 samples in 1KGP were successfully analyzed, with the final call set consisting of 36,699 MEIs from 5,675 genomes. Subfamily characterization for *Alu* MEIs and L1 MEIs was done using MELT’s CALU tool.

### Detection of L1 3’ transduction and 5’ inversion

Following the generation of a high-quality MEI call set, MELT v2.15 was used to detect L1 3’ transduction. We followed the instruction of MELT 3’ transduction identification pipeline and extracted the METRANS and MESOURCE field in the resulting VCF manually. The population frequency was calculated with the AC/AN (for offspring MEI set, we used the sum of AC and AN) and normalized across different populations.

The MELT VCF provided the position of a 5’ inversion site (from the 3’ end) through the “ISTP” field. We subtracted it from the full length of L1 (6,019 bp) to obtain the coordinate of the inversion site from the 5’ end. While comparing the inversion coordinate and the length of

L1, we removed the full-length L1 elements from the comparison set. Sites were distributed into 100 bins across the full length of L1. We compared the distribution of sequence length and inversion site position among these bins and calculated the Pearson correlation value.

### Analysis of Hardy-Weinberg equilibrium

To evaluate the genotype distributions of each MEI under the null expectations set by the Hardy-Weinberg equilibrium (HWE), we tabulated genotype distributions of autosomal MEIs per dataset and performed exact tests by “HWExactStats” function in R package HardyWeinberg v1.6.3 (Graffelman 2015). While disequilibrium may indicate disease association or population stratification, it may be the result of confusion of heterozygotes and homozygotes. We thus used the HWE test for gross quality-check of genotyping accuracy (Fig. S2), as described in (Collins et al. 2020).

### Comparison with the 1KGP MEI call set

To compare with the MEIs generated by the 1KGP (Gardner et al. 2017), we downloaded the GRCh38 version call set from the dbVar database (Lappalainen et al. 2013). Then non-reference MEIs were extracted and compared with the MEIs identified in this study, using “window” function from BEDtools v2.26.0 (Quinlan and Hall 2010). When a site was located in ±500 bp of another site, it was considered as a hit.

### Testing MELT for different genome build and joint calling

To test MELT’s performance on different genome build, we randomly generated 100 samples from the 1KGP dataset, and we got the alignment files for both GRCh37 and GRCh38 version for these samples. After which we ran MEIL v2.15 on the two dataset and filtered sites as mentioned above. Finally, we compared the results using the function “intersect” from BEDtools v2.26.0 (Quinlan and Hall 2010).

To test MELT’ performance with respect to sample size (joint calling), we randomly generated 100 samples from the NyuWa dataset and combined with the 100 random samples from the 1KGP above. Then we identified MEIs using the same pipeline as before on these 200 samples. After which we compared the call set with the MEIs detected from the 100 samples from the 1KGP with BEDtools “intersect”.

### Functional annotation

Variant Effect Predictor v99.2 (VEP) (McLaren et al. 2016) with Ensembl database version 99 (Zerbino et al. 2018) was used to annotate MEIs, with parameters “--pick --canonical --distance 1000,500”. MEIs were also intersected with enhancers from GeneHancer database (Fishilevich et al. 2017) using BEDtools v2.26.0 “intersect” function (Quinlan and Hall 2010). Only one functional consequence was kept for each MEI, and enhancers were given higher priority when a MEI was also found in non-coding genes and intergenic regions.

Mapping MEIs to the GWAS signals was done as described in a previous study (Payer et al. 2017). GWAS SNPs and their related traits were obtained from GWAS Catalog v1.0.2 (Buniello et al. 2019). We first defined the LD block region for each GWAS SNP by its proxy SNPs (r^2^ > 0.8). The LD between all the SNPs was calculated using the SNP call set generated by 1KGP phase III (The 1000 Genomes Project Consortium 2015), with plink v2.00a1LM (Chang et al. 2015). If there was no LD SNPs found in either side of the GWAS SNP, we used the median length of all predicted LD regions as the block length, centered by the target SNP. Then BEDtools v2.26.0 “intersect” function (Quinlan and Hall 2010) was employed to identify MEIs falling into these LD block regions. The complete set of these MEIs could be found in Table S5.

To qualify the enrichment of MEIs across different genomic features (Fig. 4B), we permuted 1,000 times for each MEI type with the same number as the real calls using GAT v1.3.4 (Heger et al. 2013). Each permutation set was annotated with VEP and BEDtools using the same rules as above. After counting the MEIs in each genomic feature, log2 fold changes and empirical p-values were computed. We repeated 3 times of the permutation procedure to verify the results.

### Chromosome-level analyses of MEI density

To check the distribution of MEIs throughout the genome, we used the method described by Collins *et al*. (Collins et al. 2020) and we repeated it here for clarity. Focusing on 22 autosomes, each chromosome was segmented into consecutive 100kb bins and bins overlapped with centromeres were removed. For each MEI type (*Alu*, L1, SVA and HERV-K), the number of variants in each bin was recorded to get a matrix of MEI counts per 100kb bins per autosome. To smooth the MEI counts for each MEI type, an 11-bin (∼1Mb) rolling mean per chromosome was computed. Each bin was then assigned to a percentile based on the position of that bin on its respective chromosome arm relative to the centromere. Specifically, a value of 0 corresponded to the centromere, and a value of −1 and 1 corresponded to the p-arm telomere and q-arm telomere, respectively. Finally, to compute “meta-chromosome” density shown in

Fig. 2, the normalized bin positions (i.e., −1 to 1) were cut into 500 uniform intervals, and values across all autosomes based on the normalized interval position were averaged. For the comparison of chromosome contexts (Fig. 2), normalized positions within the outermost 5% of each chromosome arm were considered as “telomeric”, the innermost 5% as “centromeric” and the other 90% of each arm as “interstitial”. Visualization of density of different MEIs on each chromosome shown in Fig. S6 was done using RIdeogram v0.2.2 (Hao et al. 2020).

### Mutation rates

Before estimating mutation rate, we exclude the MEIs failed in the HWE test (adjusted p < 0.05). MEIs in low complexity regions (Li 2014) and in reference TE sequences were also filtered, due to the inability of MELT in these regions. Watterson’s Theta (Watterson 1975) was then used to estimate the genome mutation rate of each MEI type:

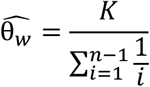

where *K*is the number of MEI site observed per MEI type in given population, and is the total number of chromosomes assessed. Then mutation rates were estimated as:

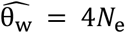

with an effective population size (i.e. *N*_*e*_) of 10,000, consistent with previous studies (Sudmant et al. 2015; Gardner et al. 2019; Collins et al. 2020). To estimate mutation rates worldwide, the average mutation rate across all five continental populations was computed, with 95% confidence interval surrounding the mean based on t distribution (Collins et al. 2020).

### SNP heterozygosity and MEI diversity

As described in a previous study (Hormozdiari et al. 2013), SNP heterozygosity was computed as the ratio of heterozygous SNPs over the length of the genome, and the mean value was used when multiple samples were considered. MEI diversity was defined as the average number of MEI differences between individuals in a population. For the NyuWa dataset (Zhang et al. 2020), high-quality SNP calls generated by the GATK v3.7 cohort pipeline (DePristo et al. 2011; Poplin et al. 2018) were used. For 1KGP3 samples, SNP calls on the human genome build GRCh38 of the were downloaded from http://ftp.1000genomes.ebi.ac.uk/vol1/ftp/data_collections/1000_genomes_project/release/20190312_biallelic_SNV_and_INDEL/. Number of heterozygous SNPs was computed by VCFtools v0.1.15 (Danecek et al. 2011) and MEI diversity by “gtcheck” function in BCFtools v1.3.1 (Danecek and McCarthy 2017).

### Database construction

We constructed the database with Bootstrap and Django. For each population, we calculated allele frequency of each MEI. All the data can be browsed in the database and downloaded from the “Download” page.

### Statistical analysis

All statistical analyses in this study were briefly described in the main text and performed using R v3.6.2 (http://CRAN.R-project.org/).

### Data Access

Complete MEI call set and other related information such as allele frequency and functional annotation are available in the companion database HMEID (available at http://bigdata.ibp.ac.cn/HMEID/).

## Acknowledgments

We thank Eugene J. Gardner for helping us in using MELT. We thank Jing Wang for valuable comments in the data analysis and critical review of the manuscript. We thank Tingrui Song for assisting the use of high-performance computing platforms. We thank the people for generously contributing samples and sequencing data to the NyuWa dataset and the 1KGP dataset. Data analysis and computing resources were supported by the Center for Big Data Research in Health (http://bigdata.ibp.ac.cn), Institute of Biophysics, Chinese Academy of Sciences. This work was supported by the National Key R&D Program of China [2016YFC0901702, 2018YFA0106901]; National Natural Science Foundation of China [31871294, 31701117, 31970647]; the 13th Five-year Informatization Plan of Chinese Academy of Sciences Grant [XXH13505-05].

## Author Contributions

T.X. and S.M.H. conceptualized and supervised the project. Y.W.N., X.Y.T., Y.R.S., Y.Y.L., Y.H.T. and Q.K. conducted data analysis. X.Y.T. built the database. Y.W.N., X.Y.T., H.X.L., P.Z. and S.M.H. drafted the manuscript, and all the primary authors reviewed, edited, and approved the manuscript.

